# Visual perceptual learning enhances functional connectivity in retinotopic space

**DOI:** 10.1101/2025.11.21.689780

**Authors:** Vikranth R. Bejjanki, Nicholas B. Turk-Browne

## Abstract

Repeated exposure to perceptual tasks improves behavioral performance. Several neural mechanisms have been proposed to account for such perceptual learning. Computational modeling suggests that plasticity in the connectivity between cortical sites may be responsible, by increasing the fidelity with which task-relevant information is transmitted through sensory hierarchies. Here we explore this theory in humans using fMRI, testing the hypothesis that perceptual learning at one location in space will increase functional connectivity between voxels in visual areas that are tuned to that retinotopic location.

Participants learned to detect one of two novel visual shape contours embedded in a noisy background in different visual quadrants. At baseline, there was no difference in behavioral sensitivity for the two shapes, nor a difference in functional connectivity between voxels in V1 and V4 responsive to the retinotopic locations of the two shapes. After training, there was a robust and selective improvement in perceptual detection of the trained shape relative to the control shape, along with increased functional connectivity between V1 and V4 voxels coding for the location of the trained shape in retinotopic space. Moreover, this increase in functional connectivity for the trained versus control shape predicted the improvement in behavioral sensitivity across participants. These results are consistent with the proposal that perceptual learning alters network dynamics so as to enhance the processing of behaviorally relevant information.

## Introduction

Repeated exposure to perceptual tasks results in large improvements in performance (Fiorentini and Berardi, 1980; Karni and Sagi, 1991; Fahle et al., 1995; Lu and Dosher, 2022). Such perceptual learning has been observed in a wide range of tasks and across modalities, suggesting that it represents a general mechanism by which humans and animals adapt their behavior to task demands (Gibson, 1963; Recanzone et al., 1992; Recanzone et al., 1993; Fine and Jacobs, 2002; Seitz and Watanabe, 2005). Notably, perceptual learning tends to be specific to the features of the trained task (Fahle and Morgan, 1996; Ahissar and Hochstein, 1997; Crist et al., 1997; Gilbert et al., 2001). For instance, behavioral improvements from training on orientation discrimination or shape identification at one location in space do not transfer to other locations or to other stimuli at the same location (Schoups et al., 2001; Lewis et al., 2009; Jehee et al., 2012; Jia et al., 2020). Such specificity suggests an early locus of learning, where neural responses retain specificity to stimulus attributes (Watanabe et al., 2002). Indeed, previous neurophysiological (Schoups et al., 2001; Yang and Maunsell, 2004; Raiguel et al., 2006; Adab and Vogels, 2011; Yan et al., 2014) and neuroimaging studies (Ress et al., 2000; Schwartz et al., 2002; Furmanski et al., 2004; Kourtzi et al., 2005; Yotsumoto et al., 2008; Bao et al., 2010; Jehee et al., 2012; Jia et al., 2020) have documented changes in neural responses and BOLD activity in early visual areas as a result of perceptual learning.

The mechanism that mediates these changes is less clear. One early and prominent proposal is that perceptual learning is mediated through a steepening––i.e., amplification or sharpening––of neural tuning curves in early sensory areas, via changes to lateral connections within these areas (Teich and Qian, 2003; Schwabe and Obermayer, 2005). This proposal is consistent with the observed changes in neural response properties in early visual areas like V1 and V4. However, neural responses are correlated *in vivo* (Zohary et al., 1994; Averbeck et al., 2006; Smith and Kohn, 2008; Ni et al., 2018), and it has been suggested that steepening of tuning curves need not lead to more informative neural representations or improved behavioral performance when noise correlations are considered (Spiridon and Gerstner, 2001; Series et al., 2004; Kohn et al., 2016). Computational modeling instead suggests that plasticity in the connectivity *between* cortical areas may contribute, by increasing the fidelity with which task-relevant information is transmitted (Bejjanki et al., 2011). Specifically, changes in connectivity between early visual areas that better match the stimulus profile — i.e., by moving closer to a matched filter for the stimulus — can induce amplification and sharpening of tuning curves in early visual areas and result in highly specific behavioral improvements.

In the current study, we test this proposal non-invasively in humans at the level of fMRI. We employed a task in which participants detected shape stimuli defined by a closed contour of collinear Gabor elements embedded in a background of randomly oriented Gabors (Fig. 1A). Two novel shapes, mirror reflections of each other, were presented in symmetric retinotopic locations in the upper-left and upper-right quadrants of visual field. We trained participants to detect one of the shapes (the “trained” shape) and measured perceptual sensitivity for this shape and the other “control” shape before and after training (Fig. 1C). We evaluated the neural consequences of this training by defining visual areas V1 and V4 retinotopically in each participant, and used population receptive field mapping (Dumoulin and Wandell, 2008; Harvey and Dumoulin, 2011) to identify voxels in these areas coding for the precise retinotopic location of the trained and control shapes. We hypothesized that, from before to after training, participants would show increased perceptual sensitivity for the trained versus control shape and greater retinotopically specific functional connectivity for the trained versus control shape; and that these behavioral and neural changes would be associated across participants.

**Figure 1.**
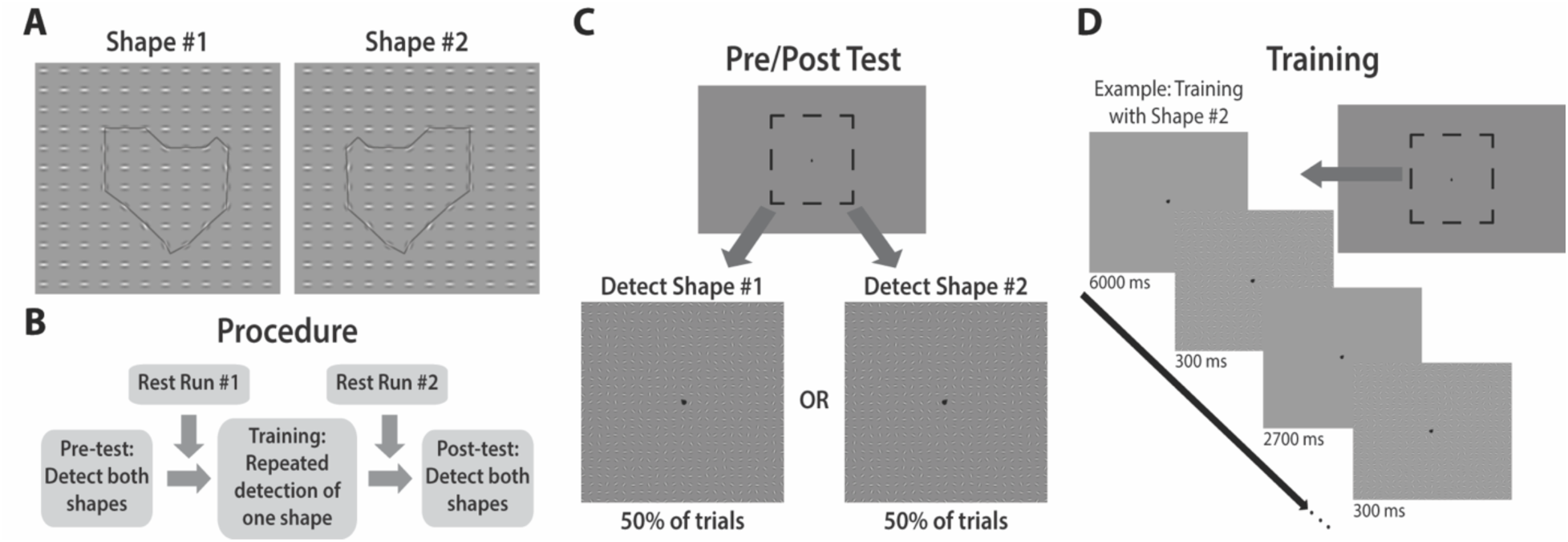
Experimental paradigm. (A) Two novel shapes were rendered with collinear Gabor elements and embedded in backgrounds of randomly oriented Gabors (horizontal background Gabors and black shape outline shown only for illustrative purposes). The two shapes were mirror reflections of each other and were presented at symmetric retinotopic locations to the upper left and upper right quadrants of fixation. (B) Participants were trained to detect one “trained” shape, with sensitivity for this shape and the “control” shape assessed before and after training. Six runs of fMRI data were collected while participants completed the training task. Rest runs were collected before and after training to assay changes in functional connectivity, during which participants fixated a central fixation symbol. (C) To measure detection behavior, participants were cued to report the presence or absence of one shape while maintaining central fixation. The cue consisted of a teardrop symbol at fixation pointing to the top-left or top-right quadrant. The dashed black box illustrates the stimulus area with the shape and background elements. No performance feedback was provided. (D) In each training trial, participants were cued to report the presence or absence of the trained shape (shape #2 in this example). Accuracy feedback was provided at the end of each training block.

## Materials and Methods

### Participants

Twenty-four naïve adults (9 males; ages: 18-31, mean = 22.8 years) with normal or corrected-to-normal visual acuity from the Princeton University community participated in two fMRI sessions for monetary compensation ($20 per hour). Four additional participants were excluded due to an inability to complete a scanning session or because they did not return for the second scanning session. This sample size is similar to, or exceeds, that used in prior fMRI studies examining perceptual learning in humans (e.g., Jehee at al., 2012; Jia et al., 2020; Kourtzi et al., 2005; Lewis et al., 2008). The Princeton University Institutional Review Board approved the study protocol, and all participants provided informed consent.

### Stimuli

The stimuli were inspired by prior studies (Kourtzi et al., 2005). Two novel shapes (Fig. 1A) were rendered with collinear Gabor elements (0.4°) and embedded in backgrounds (25 x 25 grid) of randomly oriented Gabors. Each shape involved a closed contour of eighteen collinear Gabor elements (average shape area: 4.9° x 4.9°). The two shapes were mirror reflections across the vertical meridian and were presented at symmetric retinotopic locations to the left and right of fixation in the upper visual field (Fig. 1C). The starting orientations of the background Gabor elements were drawn from a uniform distribution centered on 90° (training task: range = [38:142]; pre/post task: range = [30:150]), and the starting separation (center-to-center) of individual Gabor elements was 0.75°. To ensure that participants learned the shapes and not the background configuration, the orientation and position of each Gabor element was randomly jittered on every trial, with the jitter level set to make the shape detection task challenging (based on results from pilot tests). The display also included a teardrop shaped fixation point (0.4°), located at the center of the display (replacing the Gabor element at that location).

### Procedure

Each participant completed two experimental sessions, separated by 0-2 days. During the first session, participants completed two variants of the shape detection task: (1) training task, in which they were trained to detect one of the novel shapes; and (2) pre/post task, in which their sensitivity for detecting both shapes was assessed before and after training. Six fMRI runs were collected while participants completed the training task, preceded and followed by a rest run (Fig. 1B). The second experimental session was used to collect structural images and six additional fMRI runs for retinotopic mapping.

#### Pre/post task

During the first experimental session, participants performed the shape detection task in the MRI scanner. Stimuli were projected on a screen behind the scanner and viewed with a mirror on the head coil. Participants first performed a pre-test (Fig. 1C) to assess their baseline sensitivity for detecting the two shapes. They were told that they would be presented with several trials in which they would see a grid of small oriented objects, with a black teardrop shaped symbol in the middle. They were instructed to keep their eyes fixated on this central fixation point at all times in the experiment. On each trial of the pre-test, the teardrop pointed to the top-left or to the top-right quadrant of the display for 300 ms, after which the Gabor grid was presented for 300 ms. The display then reverted to showing a centrally oriented teardrop symbol and participants were tasked with reporting whether or not they had detected a shape in the cued quadrant while maintaining central fixation. No time limit was imposed on their response. For instance, if the fixation symbol pointed to the top-left on a particular trial, participants performed a two-alternative forced choice (2AFC) task on whether they detected the shape in the top-left quadrant. When there was a shape present, it was always the same shape in the top-left quadrant, while the other shape was always in the top-right quadrant. Participants completed a total of 80 trials during the pre-test, with 40 trials cuing the top-left quadrant and 40 trials cuing the top-right quadrant (randomly intermixed). The cued shape was present in 50% of the trials. No performance feedback was provided to participants. Trials were separated by an intertrial interval of 1 s. At the conclusion of training, participants performed a post-test identical to the pre-test.

#### Training task

One shape was designated as the “trained” shape and the other served as the “control” shape (counterbalanced across participants). The training task was almost identical to the pre-and post-tests, except that it contained only the trained shape in its same location. Thus, the teardrop always pointed to either the top-left or the top-right quadrant, depending on the participant. We collected six runs of fMRI data while participants performed the training task. Each run began with 6 s of central fixation followed by 154 trials that began every 3 s (total of 924 training trials). This timing included the stimulus display (Gabor grid with shape present on half of trials) being presented for 300 ms followed by a fixation interval of 2,700 ms. The run ended with 4 s of fixation, for a total duration of 7 m 52 s.

Participants were provided feedback on their accuracy (as percent correct) at the end of each run. During the first three training runs, the presence/absence of the trained shape was randomly ordered across trials. The order for the last three runs was mirrored with the first three runs, such that runs 1 & 6, 2 & 5, and 3 & 4 involved identical presentation orders.

#### Rest runs

We also collected fMRI data during two rest runs, each of which was the same duration as the training runs. During the rest runs, only the fixation point was presented on the screen and participants were instructed to keep their eyes open and stare at the fixation. The first rest run was collected after the pre-test but before the start of training, whereas the second rest run was collected after the last training run but before the start of the post-test (Fig. 1B). Eye position was monitored at 60 Hz with an iViewX MRI-LR eye tracker (SMI, Teltow, Germany). Due to technical difficulties, eye-tracking data could not be collected from three participants.

#### Retinotopic mapping

We performed phase-encoded retinotopic mapping during the second experimental session using standard procedures (Arcaro et al., 2009; Al-Aidroos et al., 2012). A flickering and colorful circular checkerboard (4 Hz flicker, 14° radius) was shown at fixation to stimulate visual receptive fields. During polar angle runs, which were used to map the preferred polar angle of voxels, the checkerboard was masked so that only a 45° wedge was visible, and the mask rotated clockwise or counterclockwise at a rate of 9°/s creating the perception of a rotating wedge with a 40 s period. During eccentricity runs, which were used to map the preferred eccentricity of voxels, the checkerboard was masked to create the perception of an expanding or contracting annulus. The expansion/contraction period was 40 s, and the thickness and rate of expansion of the annulus increased logarithmically with eccentricity to approximate the cortical magnification factor of early visual cortex. Participants completed six total retinotopic mapping runs (order counterbalanced across participants): one run each in which the checkerboard wedge rotated clockwise or counterclockwise, one run each in which the checkerboard ring expanded or contracted, then another run each in which the checkerboard rotated clockwise or counterclockwise. To ensure that participants were actively attending to the stimuli, they were instructed to detect when a central fixation point (radius = 0.25°) dimmed from white to gray (every 2-5 s) using a button box.

### Data analysis

#### Acquisition

Tasks were conducted with the Psychophysics Toolbox (http://psychtoolbox.org) for Matlab (Mathworks). fMRI data were acquired at the Scully Center for the Neuroscience of Mind and Behavior at Princeton University with a 3T Siemens Skyra MRI scanner using a 16-channel head coil.

Functional images for the training and rest runs were acquired using a gradient-echo, echo-planar imaging (EPI) sequence (repetition time [TR] = 500 ms; echo time [TE] = 30 ms; flip angle [FA] = 60°; matrix = 64 x 64; resolution = 3 x 3 x 3 mm; multi-band acceleration factor = 3) with 21 interleaved oblique slices (15° transverse to coronal). A short TR was used in order to prioritize collecting as many timepoints as possible for functional connectivity analyses. These parameters resulted in a partial brain volume for each participant; we positioned the slice slab over the occipital and temporal lobes to maximize spatial and temporal resolution over our *a priori* visual regions of interest (ROIs). Stimulus onsets were time-locked to the start of TRs. Functional images for retinotopic mapping runs were acquired with a gradient-echo EPI sequence (TR = 2 s; TE = 40 ms; FA = 71°; matrix = 128 x 128; resolution = 2 x 2 x 2.5 mm) with 25 interleaved slices aligned to the calcarine sulcus.

For each scanning session and functional sequence, we also collected a T1 FLASH anatomical scan, co-planar to the functional scans, to improve registration. To correct for B0-field inhomogeneity, phase and magnitude field maps were collected towards the end of each session (i.e., after all functional scans), co-planar to the functional scans and with the same resolution. Finally, a high-resolution (0.9 x 0.9 x 0.9 mm) T1 MPRAGE anatomical scan was acquired at the end of the second session, for surface reconstruction and registration.

#### Preprocessing

fMRI data were analyzed using FSL (Smith et al., 2004), FreeSurfer (Dale et al., 1999), AFNI (Cox, 1996), SUMA (http://afni.nimh.nih.gov/afni/suma/), and Matlab (Mathworks). All images were skull-stripped to improve registration. Volumes from the first 6 s of functional runs were discarded. The remaining volumes were corrected for slice-acquisition time and head-motion, high-pass filtered (retinotopic mapping runs: 100-s cutoff; training and rest runs: 128-s cutoff), spatially smoothed (retinotopic mapping runs: 3-mm FWHM; training and rest runs: 5-mm FWHM), and registered to the FLASH, MPRAGE, and Montreal Neurological Institute (MNI) standard brain.

#### Regions of interest

Based on the retinotopy runs, we manually drew occipital ROIs for V1-V4 in both hemispheres using Freesurfer, AFNI, and SUMA following previously published procedures (Arcaro et al., 2009). To visualize occipital areas simultaneously, each participant’s cortical surface was segmented along the white matter/gray matter boundary, then inflated and flattened. Functional runs from retinotopic mapping sessions were pre-whitened, corrected for hemodynamic lag (3 s), and phase-decoded to determine the angle and eccentricity of maximal stimulation for each voxel in visual cortex. We further accounted for hemodynamic lag by averaging phase estimates from clockwise/counterclockwise and expansion/contraction runs. We projected the decoded phase values onto the cortical surface and determined occipital ROIs according to the relative locations of polar-angle reversals and foveally stimulated regions (Wandell et al., 2007; Arcaro et al., 2009).

#### Resting connectivity

To infer functional connectivity between voxels in V1 and V4, we computed temporal correlations between their BOLD timeseries during the rest runs. Such correlations have been used extensively in the literature to infer the latent functional architecture at rest (Biswal et al., 1995; Fox and Raichle, 2007). After preprocessing, BOLD signal from the rest runs was scrubbed of nuisance variance using a general linear model, which contained regressors for the global mean activity, six motion correction parameters obtained from preprocessing, and the activity from two (posterior) white-matter voxels and two (posterior) ventricle voxels. We inferred the resting connectivity between pairs of voxels by computing the temporal Pearson correlation over the residuals from this model.

Because we expected learning to influence functional connectivity in a spatially specific manner, we also estimated population receptive fields (pRFs). This provided a continuous estimate of the visual field location(s) to which each voxel responded. For each ROI, we used the phase-decoded angle and eccentricity of maximal stimulation for each voxel (computed from retinotopic mapping runs), to estimate the visual field location of maximal response (Engel et al., 1997). We then combined this information — the center location (*x, y*) for the pRF — with published estimates of population receptive field size (σ) as a function of area and eccentricity (Harvey and Dumoulin, 2011), to derive an isotropic two-dimensional Gaussian pRF model of visual field locations per voxel.

We combined these estimates to compute functional connectivity between voxels in V1 and V4. We hypothesized that learning to detect a shape at one location in visual space would enhance functional connectivity between voxels in V1 and V4 that respond to that location. To test this hypothesis, we computed a weighted average of the resting connectivity between each V4 voxel in the trained and control quadrants, and all V1 voxels whose pRFs: (1) were encompassed by the pRF of the V4 voxel and (2) overlapped the trained or control shape contour. The trained/control shape contours were defined as bands spanning a width approximately equal to the average center-to-center distance between Gabors in the grid (0.75°), centered on the locations of the Gabor elements defining each shape. A V1 voxel was included if at least 30% of its pRF overlapped the shape contour. For each V4 voxel, we calculated a weighted average of functional connectivity across all included V1 voxels, with the weight determined by a Gaussian function of the distance between the V4 pRF center and the V1 pRF center. We then averaged this measure across all V4 voxels in each quadrant, and repeated this process for the two rest runs to compute estimates of functional connectivity between voxels in V1 and V4 coding for the trained and control shapes, for each participant and run.

In addition to this quadrant-averaged measure, we also sought to visualize the spatial pattern of functional connectivity before and after training. We projected the weighted average functional connectivity computed for each V4 voxel (using the above approach) back on to stimulus space, as a two-dimensional isotropic Gaussian density function, and generated an average value at each pixel in the stimulus display. Exploiting the fact that the control and trained shapes were mirror reflections of each other presented at symmetric retinotopic locations in their respective quadrants, we computed a normalized difference of the functional connectivity between V1/V4 voxels coding for the trained vs. control shape in stimulus space. Specifically, for matching polar coordinates from fixation in both quadrants, we computed the normalized difference as: [FC_Trained_ – FC_Control_ / FC_Trained_ + FC_Control_]. Because it is possible to obtain negative values for functional connectivity, and prior work has shown that the presence of negative values can bias such a metric, we computed this normalized difference after shifting all values by a constant to bring the most negative value to zero (Simmons et al., 2007).

#### Background connectivity

To assess functional connectivity between V1 and V4 voxels during the training runs, we measured background connectivity. In the presence of external stimulation, as was the case during the training runs, BOLD activity in different neural areas is correlated not only due to the connectivity between them, but also due to the synchronized stimulus-evoked responses. By exploiting the fact that evoked (stimulus-driven) and spontaneous (connectivity-driven) correlations are linearly superimposed in human fMRI data (Fox et al., 2006), the background connectivity approach to inferring functional connectivity models and linearly regresses stimulus-evoked responses out of the data, before measuring correlations in the residual spontaneous fluctuations (Al-Aidroos et al., 2012; Griffis et al., 2015; Norman-Haignere et al., 2012; Tompary et al., 2018).

After preprocessing, BOLD signal from the training runs was scrubbed of nuisance and stimulus-evoked variance using two general linear models. The first (nuisance) model contained regressors for the global mean activity, six motion correction parameters from preprocessing, and the activity from two (posterior) white-matter voxels and two (posterior) ventricle voxels. Residuals from the nuisance model were the input to the second (evoked) model. As described earlier, each training run consisted of 154 trials, with the trained shape present in half the trials, and absent in the other half. To capture the average evoked response when the shape was present/absent, we created two sets of 30 finite impulse response (FIR) regressors (one set for the shape present event, and one set for the shape absent event). Each regressor had a delta function at the corresponding TR (500 ms) following the relevant event and zeros elsewhere, resulting in a data-driven estimate of the shape and timing of the evoked hemodynamic response function (HRF) over a window of 15 s. The residuals of the FIR model, with the custom HRF removed, were correlated across V1 and V4 voxels, as described above for the rest runs, to assess background connectivity during the training runs.

#### Statistics

We used a non-parametric measure *A’* (Snodgrass and Corwin, 1988; Donaldson, 1992) to index behavioral sensitivity, as it is robust to the low hit rates we expected to obtain (particularly at the outset of the study). We used nonparametric bootstrap resampling (Efron and Tibshirani, 1986) to assess reliability for comparisons involving functional connectivity (*r*) values after variance-stabilization using Fisher’s r-to-z transformation; reported connectivity values are presented in the original *r* form to facilitate interpretation and visualization. We used the robust correlation toolbox (Pernet et al., 2013) to calculate the brain-behavior relationship across participants between the normalized difference in connectivity of voxels coding for the trained vs. control shape (formula above) and the normalized difference in behavioral sensitivity for the trained vs. control shape [*A’*_Trained_ – *A’*_Control_ / *A’*_Trained_ + *A’*_Control_]. Specifically, we used the robust skipped correlation method (accounting for bivariate outliers) to estimate the true co-efficient of association (with accurate false positive control and without loss of power) and bootstrapped 95% confidence intervals to evaluate reliability.

## Results

### Behavior

Sensitivity for detecting the target shape improved substantially across the six runs of training (repeated measures ANOVA: *F*_(5,115)_ = 5.8, *p* = 0.00008; Fig. 2A). Of the 24 participants, 20 showed numerically better performance in the last vs. first training run (Fig. 2B). Comparing performance from before to after training revealed improved sensitivity for both the trained shape (*t*_23_ = 5.88, *p* = 0.000005) and the control shape (*t*_23_ = 4.04, *p* = 0.0005), suggesting a general benefit from practicing the detection task. Critically, however, the improvement in sensitivity was greater for the trained shape than the control shape (*t*_23_ = 2.56, *p* = 0.017; Fig. 2C), consistent with specific perceptual learning of the trained shape in its quadrant.

**Figure 2.**
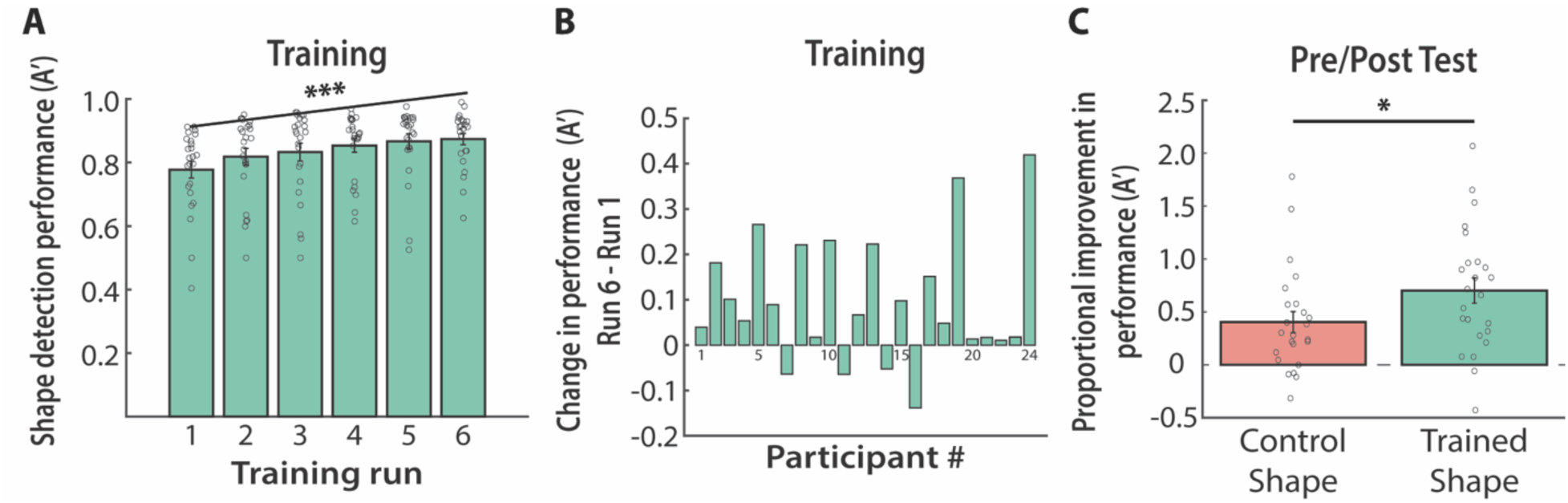
Shape detection performance, as indexed by a non-parametric measure of sensitivity (*A’*). (A) Participants improved in detecting the trained shape against a noisy background, as a function of training. (B) Most participants (20 of 24) demonstrated numerically better sensitivity for detecting the trained shape at the end of training (run 6) than at the start (run 1). (C) Shape detection improved from pre- to post-training for both the control shape and the trained shape, but significantly more for the trained shape. Dots indicate individual participants, bars represent the mean across participants, and error bars represent standard error of the mean (SEM). *** *p* < 0.001, * *p* < 0.05.

### Eye tracking

Because the trained and control shape contours were presented in the upper-left and upper-right quadrants of fixation, gaze position was monitored during the pre- and post-training rest runs to ensure that there were no systematic biases towards either quadrant. Participants reliably maintained central fixation (average horizontal displacement of gaze position from fixation did not differ from zero) during both the pre-training (*t*_20_ = 1.63, *p* = 0.12) and post-training (*t*_20_ = 0.53, *p* = 0.6) rest runs. More specifically, we confirmed that participants did not reliably differ in the extent to which their mean gaze position was deployed to the trained vs control quadrants in each run (pre-training: *t*_20_ = 0.97, *p* = 0.34; post-training: *t*_20_ = 2.0, *p* = 0.06) and that there was no significant difference between the two runs in participants’ average horizontal gaze position (*t*_20_ = 0.53, *p* = 0.6).

### Resting connectivity

What are the neural mechanisms that underlie the specific improvement we observed in shape detection performance? Here, we sought to test the theoretical proposal (Bejjanki et al., 2011) that changes in connectivity between visual areas may be responsible, by increasing the fidelity with which task-relevant information is transmitted. Specifically, we expected Gabor elements in the stimulus display to stimulate responses in the primary visual cortex, which is known to encode basic stimulus features such as orientation and spatial frequency (Hubel and Wiesel, 1968; Ferster and Miller, 2000; Nauhaus et al., 2012), and to respond robustly to Gabor-like stimuli. Furthermore, we expected the representation of the target shape to arise further downstream in V4, which receives input from the primary visual cortex, has large receptive fields (thereby integrating input information pertaining to multiple Gabor elements), and is known to encode shape and object information (Kastner et al., 2000; Thielscher et al., 2008). Thus, we expected the stimulus displays used in our task to be represented by retinotopically organized voxels in visual areas V1 and V4 (Wallis and Rolls, 1997; Riesenhuber and Poggio, 2000). We hypothesized that perceptual learning of a shape at one location in visual space would strengthen the coupling between voxels in V1 and V4 with overlapping population receptive fields tuned to that location.

To test this hypothesis, we compared V1-V4 functional connectivity during the pre- and post-training rest runs in the visual field quadrants where the trained and control shapes appeared (Fig. 3A). Because the rest runs occurred immediately before and after training, these runs provided a clean measure of how baseline functional connectivity was altered by perceptual learning. In the pre-training rest run, we did not observe a baseline difference in functional connectivity between V1 and V4 voxels coding for the quadrant of the trained vs. control shape (bootstrap *p* = 0.35). However, in the post-training rest run, this difference was marginally significant (*p* = 0.06); the interaction of run by condition was not significant (*p* = 0.11).

**Figure 3.**
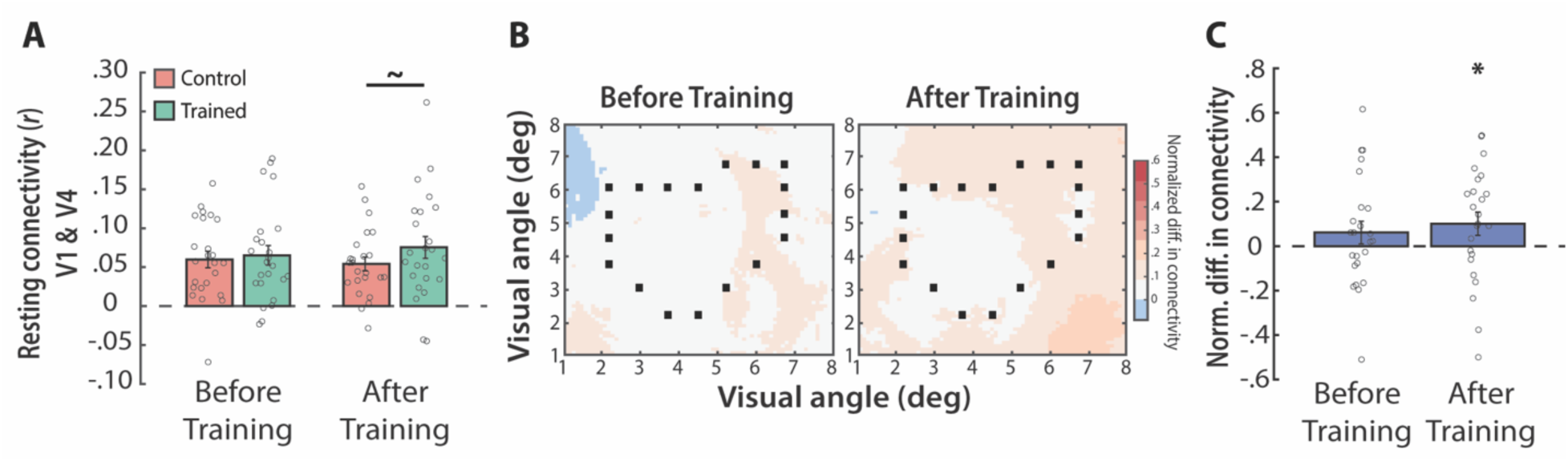
Resting functional connectivity before and after training. (A) Weighted average of the resting connectivity between voxels in V1 and V4 coding for the quadrant of the trained (green) and control (pink) shapes. Dots indicate individual participants, bars represent the mean across participants, and error bars represent SEM. (B) Group visualization in stimulus space of the difference in resting connectivity between V1 and V4 voxels coding for the locations of the trained vs. control shape. The normalized difference in V1-V4 connectivity, averaged across participants, is shown for each cell in a 7°x7° grid centered on the shape contour. The locations of the individual Gabor elements that made up the shape contour are depicted with black squares. (C) Normalized difference in functional connectivity, specific to the shape contour, before and after training. ∼ *p* = 0.06, * *p* < 0.05.

This analysis included V1 and V4 voxels whose preferred retinotopic locations were anywhere in the trained and control quadrants, which may have diluted effects that were retinotopically specific to the precise location of the trained shape contour. We therefore examined the spatial distribution of differences in functional connectivity by projecting the functional connectivity of each V4 voxel (weighted average of connectivity with overlapping V1 voxels) onto stimulus space and averaging across values at each pixel (Fig. 3B). To quantify contour-specific changes in functional connectivity, we computed a normalized difference in functional connectivity along the contour of the trained vs. control shape in the pre- and post-training rest runs (Fig. 3C). Consistent with the quadrant-level results, there was no reliable difference in the pre-training run (*p* = 0.11), but importantly, the effect in the post-training run was clearer when restricted to the shape contour (*p* = 0.03).

### Brain-behavior relationship

These results are based on averaging across all participants. However, there was considerable variability in behavioral perceptual learning (Fig. 2B). We would not expect, for example, the four participants who did not improve behaviorally during training to exhibit greater functional connectivity for the trained shape from pre- to post-training. Indeed, when we restricted the analysis shown in Fig. 3A to the 20 participants who had numerically greater behavioral sensitivity in the last vs. first training run (Fig. 4A), the post-training difference in V1-V4 functional connectivity between trained vs. control quadrants changed from marginal to significant (*p* = 0.009) despite the smaller sample size; the run by condition interaction was marginal (*p* = 0.05). As expected, there was still no pre-training difference (*p* = 0.29).

**Figure 4.**
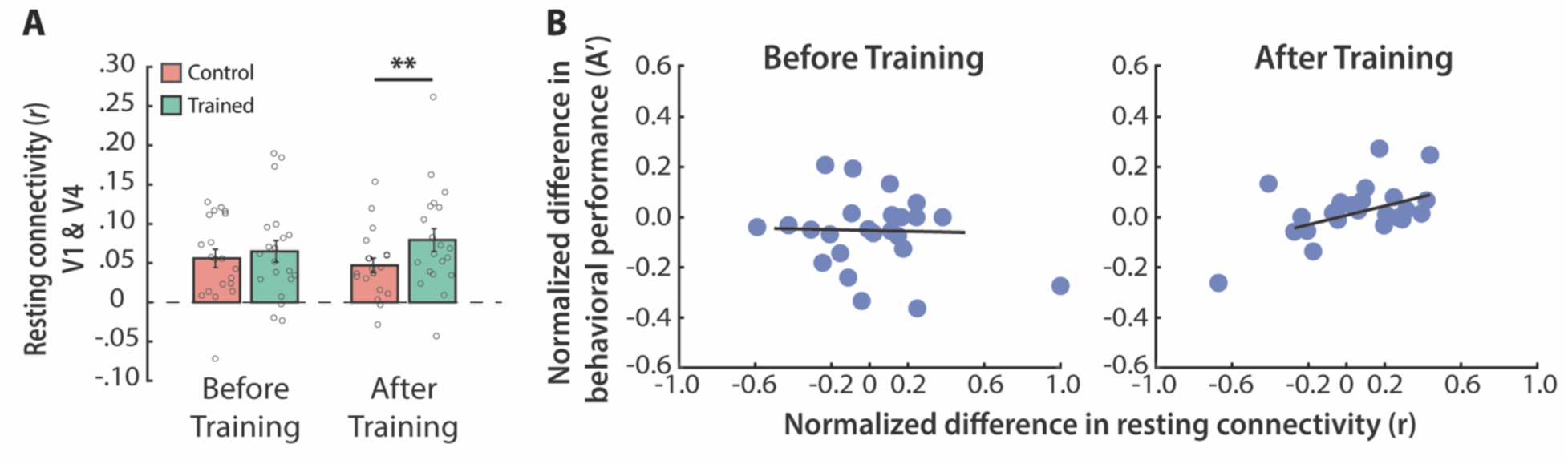
Brain-behavior relationships. (A) Functional connectivity, before and after training, for the 20 (of 24) participants who showed improved behavioral performance in the last vs. first training run. Dots indicate individual participants, bars represent the mean across participants, and error bars represent SEM. (B) Robust correlations between the normalized difference in resting functional connectivity (FC) for voxels coding for the quadrant of the trained vs. control shape and the normalized difference in behavioral sensitivity for detecting the trained vs. control shape. Dots represent individual participants.

This binary approach to identifying learners vs. non-learners obscures continuous variance across individuals. We thus performed a quantitative analysis of the relationship between learning-induced changes (normalized difference of trained vs. control shapes) in behavioral sensitivity and resting functional connectivity across the full sample (Fig. 4B). Indeed, there was a significant positive brain-behavior correlation after training (*r_skipped_* = 0.55, bootstrapped 95% CI [0.12, 0.75]). This was not a generic effect of individual differences, as the relationship was not reliable before training (*r_skipped_* =-0.02, bootstrapped 95% CI [-0.39, 0.40]). Thus, for a given participant, the extent to which functional connectivity between V1 and V4 voxels coding for the trained (vs. control) shape increased with training predicted the extent to which behavioral sensitivity increased.

### Functional connectivity during training

We included rest runs immediately before and after training in order to obtain a clean primary measure of learning-induced changes in functional connectivity between visual areas unconfounded by stimulus responses and task demands. For completeness, we also explored a secondary measure of changes in functional connectivity *during* training. Namely, we calculated background connectivity between voxels in V1 and V4 by first regressing out noise sources and stimulus-evoked responses, and then computing correlations in the residual BOLD timeseries. Although the background connectivity between V1 and V4 voxels coding for the quadrant of the trained shape was numerically greater than that between voxels coding for the quadrant of the control shape, these differences were not reliable (all *p*s > 0.1; Fig. 5A).

**Figure 5.**
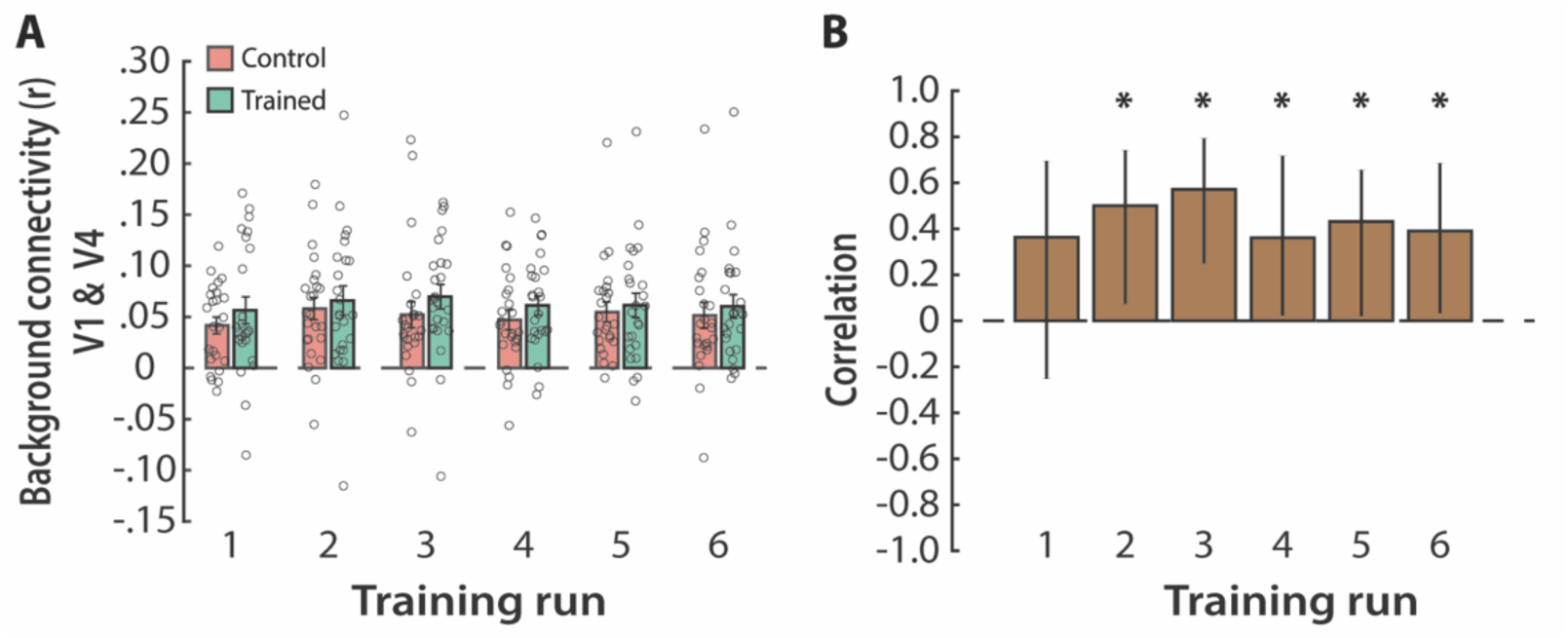
Functional connectivity during training. (A) Weighted average of the background connectivity between voxels in V1 and V4 coding for the quadrant of the trained (green) and control (pink) shapes. Dots indicate individual participants, bars represent the mean across participants, and error bars represent SEM. (B) Robust correlations between the normalized difference in background connectivity for voxels coding for the quadrant of the trained vs. control shape and the normalized difference in post-training behavioral sensitivity for the trained vs. control shape. Bars represent best-fit correlation coefficients and error bars represent 95% bootstrapped confidence intervals, obtained using the robust skipped correlation method. * *p* < 0.05.

Despite the lack of an overall effect, we next explored whether background connectivity was behaviorally relevant. For each training run, we correlated the trained vs. control difference in V1-V4 background connectivity with the difference in behavioral sensitivity for the trained vs. control shape after training. During the first training run, there was no relationship between background connectivity and post-training behavioral sensitivity (*r_skipped_*=0.36, bootstrapped 95% CI [-0.25,0.69]). However, during each of the remaining five runs, the relationship was positive and reliable (Run 2: *r_skipped_*=0.5, bootstrapped 95% CI [0.07,0.74]; Run 3: *r_skipped_*=0.57, bootstrapped 95% CI [0.25,0.79]; Run 4: *r_skipped_*=0.36, bootstrapped 95% CI [0.02,0.61]; Run 5: *r_skipped_*=0.43, bootstrapped 95% CI [0.02,0.66]; Run 6: *r_skipped_*=0.39, bootstrapped 95% CI [0.03,0.69]; Fig. 5B). These results suggest that behaviorally relevant changes in functional connectivity occurred relatively early in training, providing a more granular view of the timecourse of learning than the before-and-after snapshots obtained from rest runs.

## Discussion

Repeated exposure to perceptual tasks improves behavioral performance. The current study examined the neural basis for such perceptual learning in the human brain. The specificity of perceptual learning suggests that it occurs at an early stage of visual processing, consistent with changes in early sensory areas observed with neurophysiology (Schoups et al., 2001; Yang and Maunsell, 2004; Raiguel et al., 2006; Adab and Vogels, 2011; Yan et al., 2014) and neuroimaging (Ress et al., 2000; Schwartz et al., 2002; Furmanski et al., 2004; Kourtzi et al., 2005; Yotsumoto et al., 2008; Bao et al., 2010; Jehee et al., 2012; Jia et al., 2020). The mechanism driving these changes is often assumed to be steepening (i.e., amplification or sharpening) of neuronal tuning curves in early sensory areas (e.g., Teich and Qian, 2003; Schwabe and Obermayer, 2005). Here we tested an alternative mechanism whereby the connectivity between visual areas comes to more closely match the stimulus profile, improving the transmission of task-relevant information (Bejjanki et al., 2011).

Our findings provide evidence consistent with this matched filter hypothesis. After substantial exposure to a shape contour in one visual quadrant, there was robust and specific improvement in behavioral sensitivity to that trained shape relative to a mirrored control shape in the opposite quadrant. This was accompanied by an increase in resting functional connectivity between V1 and V4 voxels whose population receptive fields overlap the trained shape contour in retinotopic space. Furthermore, this neural change predicted the behavioral improvement across participants. Finally, we demonstrated that the enhanced functional connectivity between V1 and V4 voxels coding for the trained shape (relative to those coding for the control shape) begins to take on behavioral significance, in terms of predicting eventual differences in post-training performance, early in training. These findings do not refute other accounts of perceptual learning in the brain and may be compatible with the existence of multiple mechanisms. However, they do provide positive evidence for learning-induced plasticity in the connectivity between visual areas.

The current study may also provide a new perspective on past findings. For instance, Jehee et al. (2012) found that extensive practice on an orientation discrimination task resulted in selective enhancement of the neural representation of trained orientations in primary visual cortex. The authors speculated that their results might reflect a sharpening of the underlying population response, a reduction in noise correlations, or some combination. Our findings suggest that this could have been mediated by enhanced functional connectivity between the LGN and V1. As another example, Lewis et al. (2009) showed that perceptual learning can modulate resting connectivity in broad functional networks across the human brain in a task-specific manner. Our findings build upon this work by showing that perceptual learning can modulate fine-grained, retinotopically specific functional connectivity between early sensory areas to better match the stimulus profile.

Future work could explore the extent to which the specific parameters of our study influenced the results. For instance, we utilized a task that has been previously shown to engender neural changes in early sensory areas (Kourtzi et al., 2003; Kourtzi et al., 2005). Whether perceptual learning involves early or late areas (or both) is likely to depend on the difficulty of the task, the sensory modalities involved, and the nature of feedback and training received by participants (Chowdhury and DeAngelis, 2008; Law and Gold, 2008; Kahnt et al., 2011; Diaz et al., 2017). For example, Kourtzi et al. (2005) failed to observe changes in early sensory areas when shape contours were presented in a “high-salience” configuration (background Gabors were uniformly oriented such that the target shape popped out); instead, changes occurred in later areas such as object-selective lateral occipital cortex.

Ultimately, the theoretical perspective that inspired our study (Bejjanki et al., 2011) could apply to perceptual learning in both earlier and later stages of processing, with changes in the connectivity in the corresponding brain regions improving the transmission of task-relevant information to subsequent stages. Indeed, Jia et al. (2020) used ultra-high-field laminar fMRI to show that extensive training on visual orientation discrimination leads to enhanced orientation-specific representations in primary visual cortex, as well as enhanced feedforward connectivity between early sensory areas and subsequent task-relevant decision areas. The authors concluded that this pattern is consistent with task-relevant re-weighting of feedforward connectivity, providing further support for the matched filter hypothesis.

Another notable feature of our study is that it involved only one hour (924 trials) of training, within a single experimental session. Previous studies have documented neural and behavioral changes after a wide range of training durations, from single-shot to several days (Ahissar and Hochstein, 1997; Fine and Jacobs, 2002). Notably, although many studies have documented increased BOLD activation in early sensory areas as a result of perceptual learning, this effect appears to vanish after extensive practice, despite the persistence of behavioral improvements (Yotsumoto et al., 2008) and enhanced neural representations of task-relevant information (Jehee et al., 2012). Future work could examine the influence of more extended practice on task-relevant changes in resting functional connectivity (see Kang et al., 2018). On a related note, our results do not speak to the durability of changes in resting connectivity beyond the conclusion of the study. A future study could explore these changes over time and how their durability relates to the duration of training.

In conclusion, human observers have a remarkable capacity to selectively adapt their perceptual behavior in response to task demands. Here, we provide evidence that such perceptual learning is mediated, at least in part, by changes in network dynamics in the human brain that enhance the fidelity with which task-relevant information is transmitted to improve probabilistic inference.

## Acknowledgements

We are grateful to M. Arcaro for assistance with retinotopic mapping.

## Funding Information

This work was supported by NIH R01 EY021755 and the Canadian Institute for Advanced Research.

## Notes

### Competing Interest Statement

The authors have declared no competing interest.

